## Supplementary figures and images for "The triad interaction of ULK1, ATG13, and FIP200 is required for ULK complex formation and autophagy"

### Supplementary Figures

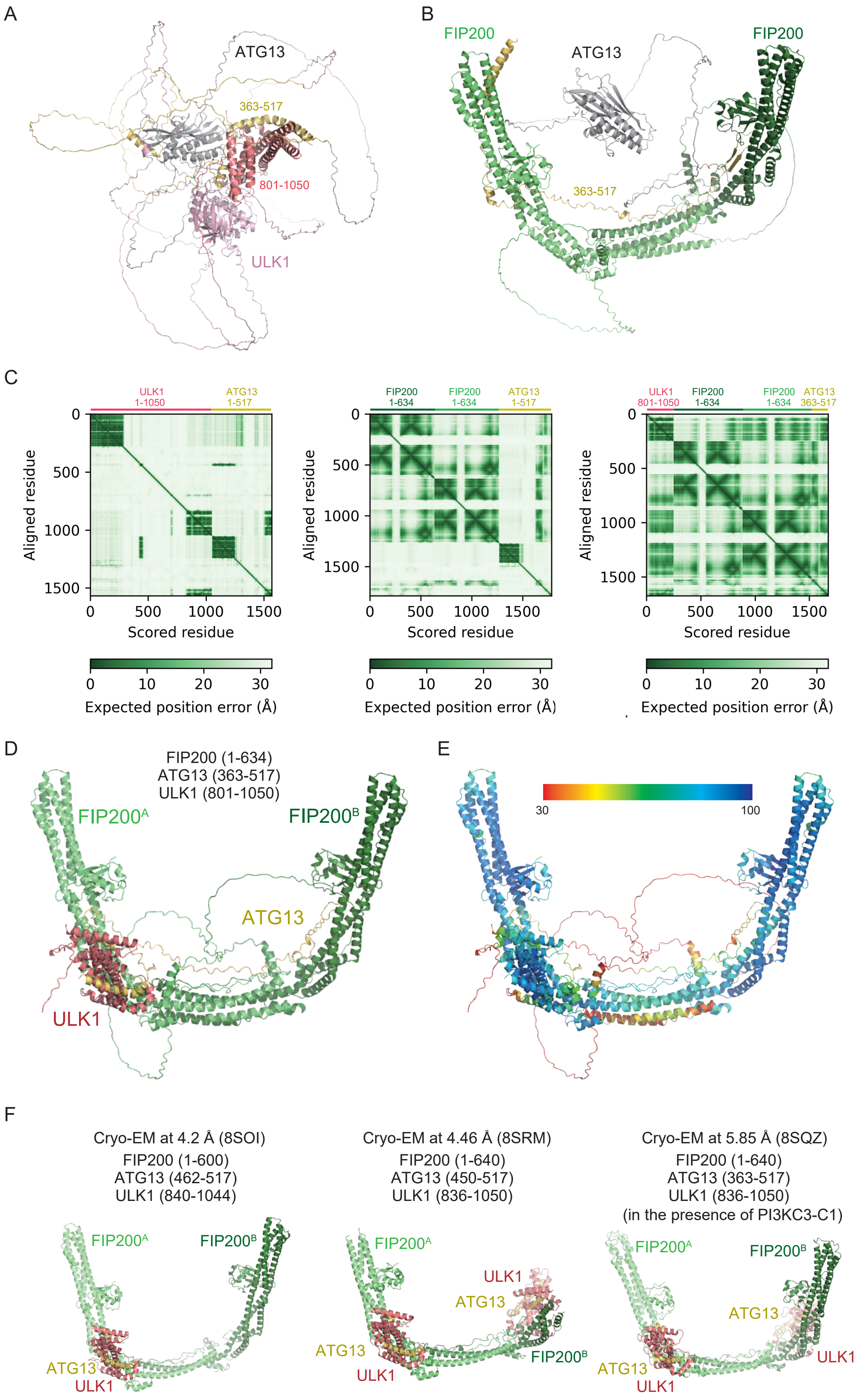

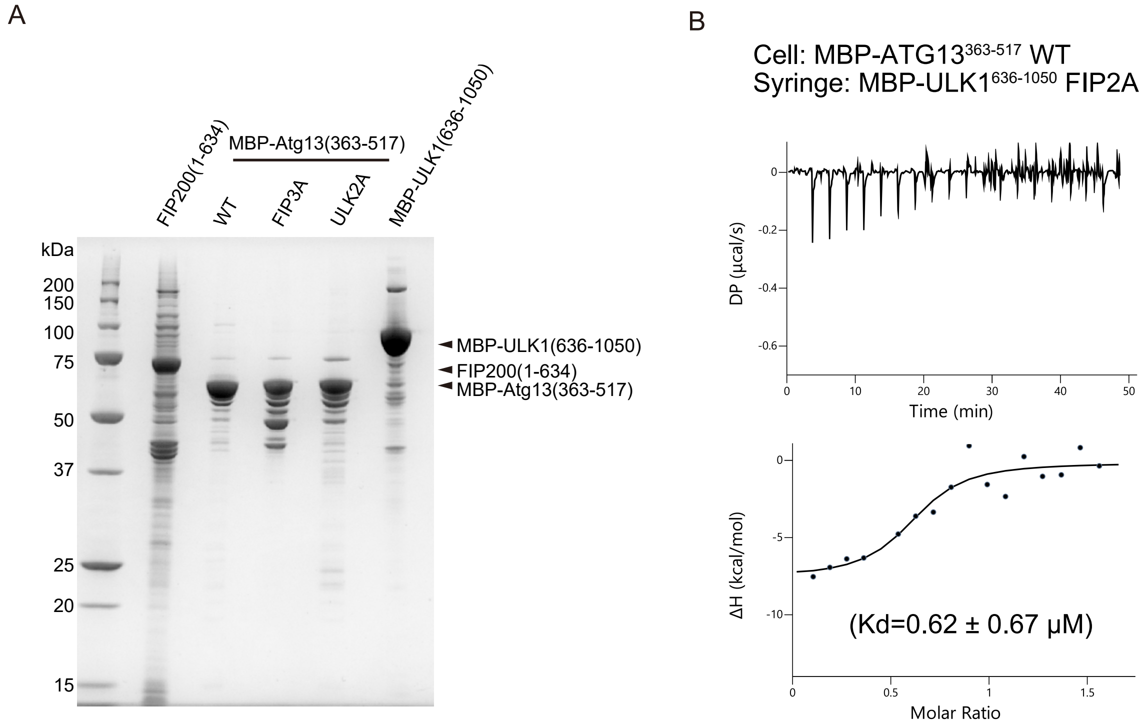

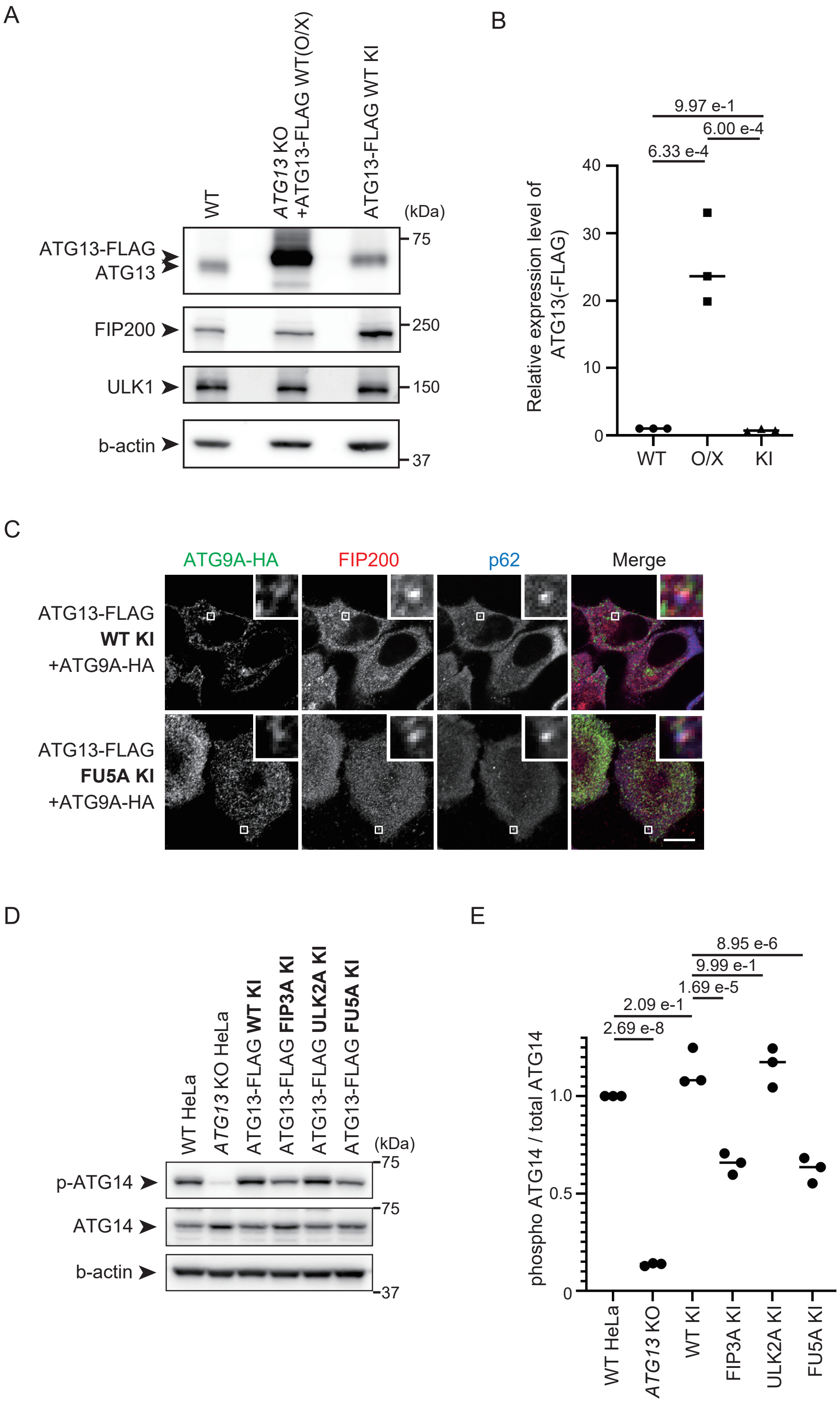
